# Exploring bacterial diversity via a curated and searchable snapshot of archived DNA sequences

**DOI:** 10.1101/2021.03.02.433662

**Authors:** Grace A. Blackwell, Martin Hunt, Kerri M. Malone, Leandro Lima, Gal Horesh, Blaise T.F. Alako, Nicholas R Thomson, Zamin Iqbal

## Abstract

The open sharing of genomic data provides an incredibly rich resource for the study of bacterial evolution and function, and even anthropogenic activities such as the widespread use of antimicrobials. Whilst these archives are rich in data, considerable processing is required before biological questions can be addressed. Here, we assembled and characterised 661,405 bacterial genomes using a uniform standardised approach, retrieved from the European Nucleotide Archive (ENA) in November of 2018. A searchable COBS index has been produced, facilitating the easy interrogation of the entire dataset for a specific gene or mutation. Additional MinHash and pp-sketch indices support genome-wide comparisons and estimations of genomic distance. An analysis on this scale revealed the uneven species composition in the ENA/public databases, with just 20 of the total 2,336 species making up 90% of the genomes. The over-represented species tend to be acute/common human pathogens. This aligns with research priorities at different levels from individuals with targeted but focused research questions, areas of focus for the funding bodies or national public health agencies, to those identified globally as priority pathogens by the WHO for their resistance to front and last line antimicrobials. Understanding the actual and potential biases in bacterial diversity depicted in this snapshot, and hence within the data being submitted to the public sequencing archives, is essential if we are to target and fill gaps in our understanding of the bacterial kingdom.

## INTRODUCTION

The widespread availability of high-throughput sequencing has resulted in a huge wealth of bacterial genomic data collected from countries all over the world that are shared openly through the public archives, representing a unique and essential resource. Studying the extreme diversity of bacterial species is of broad interest to communities with focuses of basic science, agriculture and medicine. Beyond their primary function of genomic data storage, sequence repositories show trends in funding, biases in the collection strategies of bacteria and even reveal the drive and focus of individuals pursuing particular lines of research. Sequence read data is held by members of the International Nucleotide Sequence Database Collaboration (INSDC) (1), who include DNA Data Bank of Japan (DDBJ), European Bioinformatics Institute (EMBL-EBI) and National Centre for Biotechnology Information (NCBI). Submission of genomic data to the ENA (EMBL-EBI) or its INSDC partners (DRA for DDBJ, SRA for NCBI) has become a central and mandatory step in dissemination of research to the scientific community and a way to ensure open and free access to data (1). Each of these repositories host the raw read data as well as genome assemblies, at different levels of completeness, that have been submitted by a user. These archives are continuing to grow at a remarkable rate with current estimation of doubling time of datasets in the ENA to be just over 2 years (https://www.ebi.ac.uk/ena/browser/about/statistics). The ever-increasing data size presents difficulties for storage capacity. Even more, a general user’s ability to access and effectively use the data is restricted, whether due to their computational skills, the biological question, the volume of data, the IT infrastructure or other resources required. The capacity to effectively and quickly identify datasets relevant to a user is a significant challenge, and currently DNA searches are not supported across all datasets. Furthermore, once a user has their list of datasets, significant processing for quality control and extraction of relevant data is required prior to applying specific analyses. Over time, many of these processing steps will be performed repeatedly by different researchers worldwide.

Other databases exist that provide a higher level of curation, including NCBI’s Refseq (2). Refseq (195,316 assemblies in September 2020) is composed of a selection of assemblies that have been submitted to INSDC databases that meet their quality control requirements, and most have been re-annotated using NCBI’s prokaryotic genome annotation pipeline (3) to provide consistency across the data. The assemblies are widely used for taxonomic identification (4, 5), but are also commonly used to examine the distribution of genes or elements of interest, or as test sets for new algorithms or programs (6, 7). However, the Refseq assemblies have been collated progressively over time using a range of sequencing technologies and assembly algorithms, making the assemblies less consistent and so potentially more problematic for drawing wide-ranging conclusions (8, 9).

Attempts to standardise the assembled dataset tend to have a community focus such as Enterobase which holds sequencing data from the *Enterobacteriaceae*, and includes curated genome data for 466,670 *Salmonella, Escherichia/Shigella, Clostridioides, Vibrio, Helicobacter, Yersinia* and *Moraxella* genomes (10). Enterobase gathers sequence data with associated metadata by actively searching for new sequence submissions for supported genera or through direct submissions. The raw data is then processed in a uniform way (assembly and annotation) and basic organism-specific typing is performed (10). However, whilst standardised, the scope of this type of database is by definition limited. Depending on an individual’s focus this can act to further fragment genome data and lead to even more incompatibility issues if the complete genome dataset, agnostic of organism, is to be analysed.

Here, we present a uniformly processed archive of 661K bacterial genomes that were available in the ENA at the end of November in 2018. Through the quality control steps, characterisation of the assemblies and the provision of a searchable database we remove some of the technical barriers for the interrogation of the public sequences. We use this data to examine the composition of the sequencing archives and in doing so highlight the influence of sampling and sequencing trends on the composition of these public databases.

## RESULTS

### Construction of a unified resource

On the 26^th^ of November of 2018 there were 880,947 bacterial read sets available in the ENA. Those that were single-ended or were sequenced on the PacBio or nanopore platform were removed, and 710,696 unique sample IDs were submitted to an assembly pipeline (see methods), yielding 664,877 assemblies. A subset of these (3,472 assemblies) had a genome length significantly outside that expected of a bacterial organism (smaller than 100 Kb or larger than 15Mb), leaving 661,405 standardised assemblies. Quality control and general characterisation were performed on these 661K assemblies (see methods). Standard quality control cut-offs, many of which are consistent with the threshold for inclusion for Refseq, were applied to identify genomes that were of high assembly quality. These assemblies represent complete or almost complete genomes that weren’t overly fragmented and had a genome length within an acceptable tolerance (+/- 50%) of that expected of its species. 639,981 assemblies reached or exceeded these thresholds (Supplementary Figure 1A, filter status 4).

Using Kraken2 and then refining the output using Bracken, it was evident that of the read sets contributing to these assemblies 94.1% (602,406/639,981) showed the major taxonomic species to account for 90% or greater of the total reads in that read set (Supplementary Figure 1C). Hence, there was little evidence of mixed samples or significant contamination. Importantly, lowest common ancestor approaches are not ideal if the major taxa is a member of a species complex. Therefore, we calculated an adjusted abundance (see methods) for members of the *Mycobacterium tuberculosis* complex, *Bacillus cereus* sensu lato group, or where genera or species represent taxonomic anomalies such as the division of *Shigella* sp. and *Escherichia coli* which is based on clinical imperative rather than a true taxonomic distinction (11, 12). For some species, including *Burkholderia pseudomallei, Bordetella pertussis, Mycobacterium ulcerans* and *Campylobacter helveticus*, the major species abundance in more than 97.6% of their assemblies were less than 90% using these approaches (Supplementary Figure 2), despite passing earlier quality control thresholds for contamination (Supplementary Figure 1D). This indicates that there are likely limitations with the methods for species identification used here. Of note, 89.8% (593,628) of the assemblies in the 661K had been submitted with species metadata that was consistent with the major species we identified *in silico* from sequence.

To facilitate access and usage we have added three indices that can be downloaded along with the 661,405 assemblies. The COBS (13) index allows the user to search for single nucleotide variations and polymorphisms, as well as whole genes or even extrachromosomal elements such as plasmids. Secondly the Minhash index (14), containing signatures of the assemblies can be used to search for matches to any query genomes (*i.e*. to find similar genomes). A third index, constructed using the library sketching function of PopPunk (15), includes the calculated core and accessory distances between the 661K assemblies. Genetic distance estimations for any subset of assemblies can be extracted quickly and easily from this index.

### Diversity and sequencing trends

The 639,981 high-quality assembled genomes comprised 2,336 species (Supplementary Figure 1B), and the breakdown of the genomes based on the year that they were made public in the ENA is shown in Supplementary Figure 3A. Despite the considerable number of species in this dataset, sampling was extremely unevenly distributed, with just 20 species accounting for 90.6% of the assembled data set (Figure 1A). Within this, *Salmonella enterica* accounted for almost a third of the data (28.0%), while *E. coli* (13.4%), *Streptococcus pneumoniae* (7.9%), *Staphylococcus aureus* (7.4%) and *M. tuberculosis* (7.3%) combined constituted over 35% of the remaining assemblies (Figure 1A). The final 9.4% of the assemblies comprised 2,315 species *i.e*. 99.1% of the species diversity, of which 1,861 species contributed to just 1% of the total submitted and processed data (Figure 1B). A similar trend is revealed when the contributing sequencing projects are examined, with 50% of the data originating from 50 sequencing projects (Supplementary Figure 3B), a small fraction of the total 23,316 projects. The majority of the sequencing projects (20,002) only yielded a single assembly. Unsurprisingly, three of the five largest projects focus on *S. enterica*. These include the PulseNet *S. enterica* genome sequencing project (PRJNA230403, 59,011 assemblies, 2014 onwards) run by the Centre for Disease Control (16), the Salmonella Reference Service (Gastrointestinal Bacteria Reference Unit) from Public Health England (PRJNA248792, 35,942 assemblies, 2014 onwards) (17) and the GenomeTrakr project (PRJNA186035, 19,418 assemblies, 2012 onwards) run by the US Food and Drug Administration Center for Food Safety and Applied Nutrition (18). The ramping up of these large public genomic surveillance projects in 2014 contributed to *S. enterica* dominating as the major bacterium sequenced from 2015 (Figure 1C, Supplementary Figure 3C). The Global Pneumococcal Sequencing GPS study I (PRJEB3084, 20,667 assemblies), which focuses on *S. pneumoniae* (19, 20), and a US public health project focusing on *E. coli* and *Shigella* (PRJNA218110, 20,508 assemblies, 2014 onwards) (16) are the 3^rd^ and 4^th^ largest projects in the archive. Specific interests of individuals or groups have also contributed to these sequencing trends, though the impact is more obvious in the earlier years, where organisms such as *Bordetella pertussis* (PRJEB2274) (1) and *Salmonella bongori* (PRJEB2272) (2) were prominent but were overshadowed in later years (Figure 1C).

**Figure 1.**
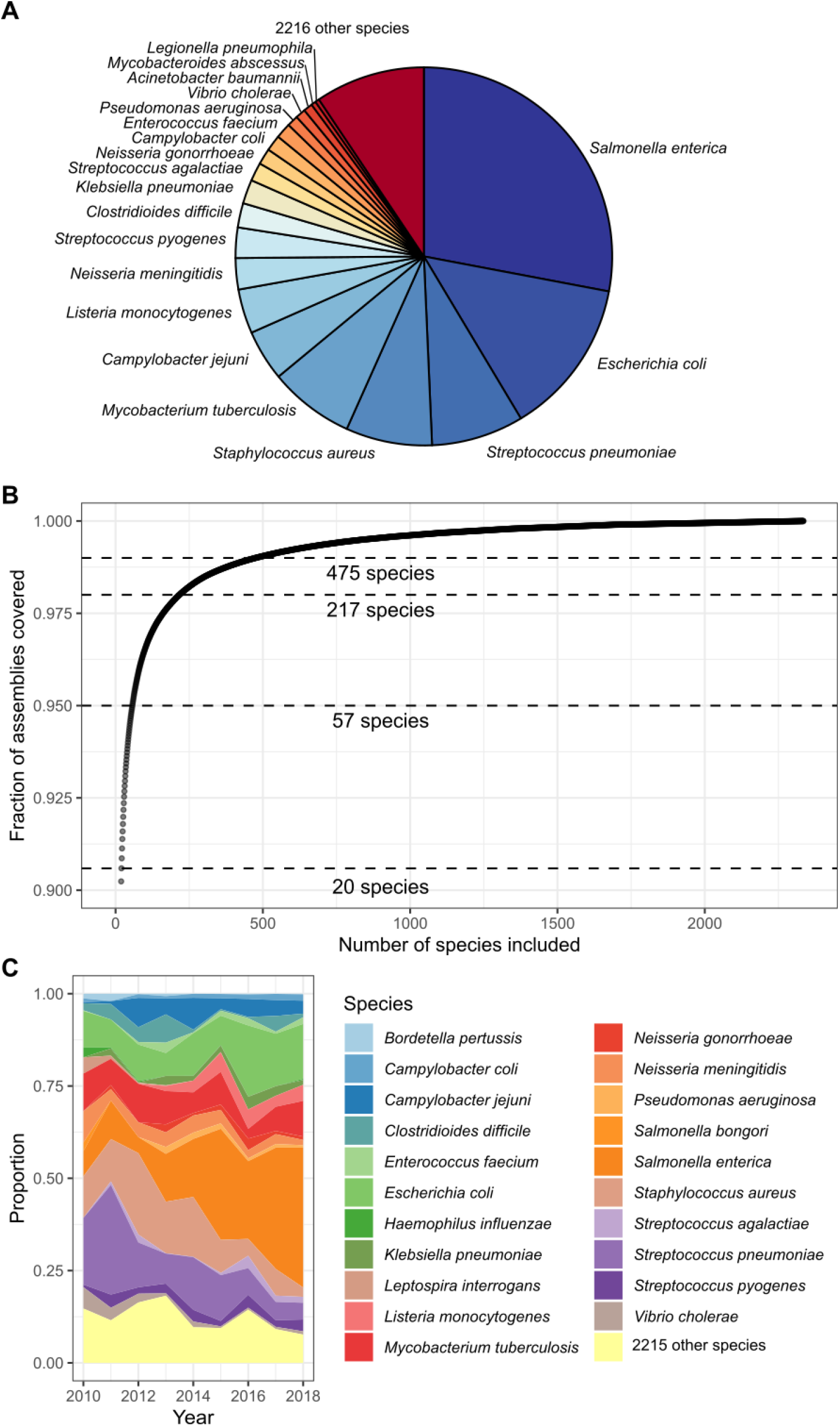
Species composition of the 639,981 high-quality assemblies. A) Relative proportions of species to the data as a pie chart. Note that 90% of the assemblies are from 20 bacterial species. B) Fraction of assemblies covered by accumulating bacterial species. C) Tracking proportions of the top 10 bacterial species for each year.

### Distribution of and accumulation of antimicrobial resistance genes

One of the major selective forces that has perturbed bacterial populations has been the development and wide-spread therapeutic use of antimicrobials since the 1940’s (21–23). Antimicrobial resistance (AMR) is highlighted as one of the greatest threats to human health (24, 25). It has been estimated that if no action is taken, 10 million people worldwide could die from drug resistant infections each year by 2050 (26). We have genotypically predicted the presence of AMR, virulence and stress response genes for all assembled genomes (see methods), but the results shown below are for the 602,407 high quality genomes with a confident major species (>90% abundance major species), unless specified otherwise. Our approach detects both genes that are core to a species, usually located on the chromosome(s), as well as those which have been horizontally acquired and are chromosomally located or otherwise located in extrachromosomal elements, such as plasmids. However, specific point mutations/deletions are not considered in this analysis.

In total, 1,655 known AMR gene variants were identified. Gene variants showed different distribution ranges across the assembled taxa with 135 gene variants detected in two or more phyla. This reduced to just 73 when a stricter 98% threshold for abundance of the major species was set to limit the effects of low level contamination commonly seen in submitted data (Supplementary Figure 4). Gene variants with more restricted distribution patterns, such as those found only within a particular genus or species could represent variants that have recently arisen within that population, or were restricted directly, through for example gene expression, or indirectly based on the host range of the plasmid or vector that carries them. For example the distribution patterns of the colistin resistance genes, first identified in 2016 (27), are at most detected within a bacterial order (*mcr-9*), or more commonly within a class (eg. *mcr-1, mcr-3, mcr-5*), while some are only present in a single species (*mcr-1.7, mcr-4.1*).

An important trend seen in our data is the relative number of genomes carrying multiple AMR genes. The count of AMR genes in each genome for two of the most represented orders - Bacilli and Gammaproteobacteria - are shown in Figure 2. Most genera within the Bacilli contain genomes with fewer than 10 antimicrobial resistance genes. Some genomes belonging to *Bacillus* and *Streptococcus* possess up to 10 or 11 resistance genes, while those from *Enterococcus* and *Staphylococcus* can carry up to 23 and 25 resistance genes in a single genome, respectively (Figure 2A). It’s important to note that some of these resistance genes are core to a species (genes found in >95% of the genomes belonging to that species). For example, 3 of the genes counted in *Enterococcus* (*aac*(6’)-*Ii, msrC* and *eatA*) were core, consistent with previous analysis (28, 29). Similarly in *S. aureus*, the *tet38* efflux pump (30) is a core gene.

**Figure 2.**
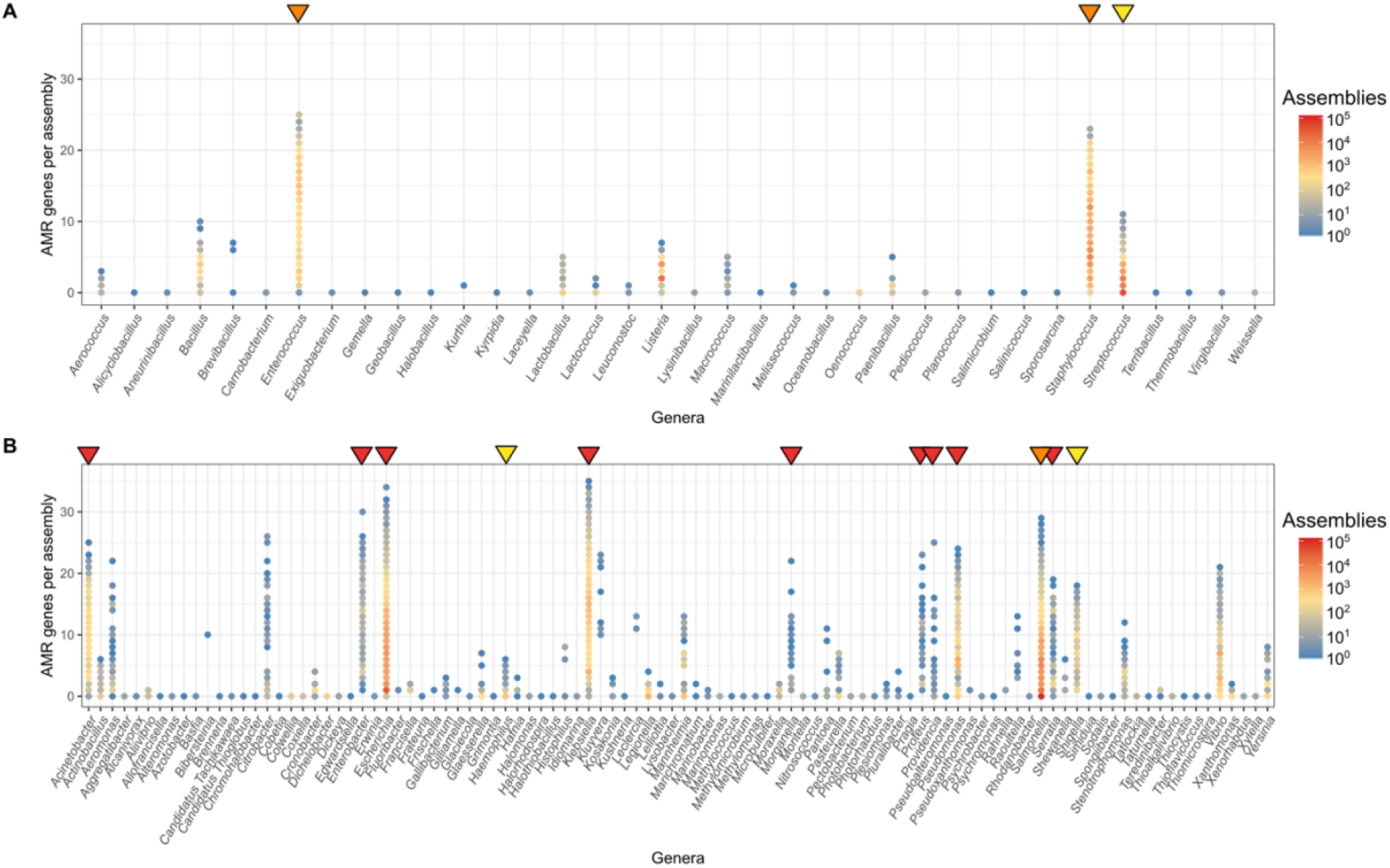
Number of AMR genes in individual genomes of the orders A) Bacilli and B) Gammaproteobacteria. Arrows above indicate genera that contain species that have been determined by the WHO to be of critical (red), high (orange) and medium (yellow) priority pathogens for research and development into new antibiotics (24).

Gammaproteobacteria represent a large proportion of the Gram-negative pathogens with many of the genera in this class possessing high AMR gene counts (Figure 2B). Most notably, *Acinetobacter, Escherichia, Klebsiella, Pseudomonas* and *Salmonella* with a small number of *E. coli* and *K. pneumoniae* genomes containing over 30 different AMR genes concurrently, while only 1 and 4 genes of these were species core genes, respectively.

The above genera with high AMR gene carriage (Figure 2) harbor species identified by the WHO as priority pathogens for research and development into new antibiotics (24). The different categories described by the WHO (critical, high and medium) are displayed in Figure 2, using the red, orange and yellow triangles. Other genera, not on the WHO priority list, show a high abundance of antimicrobial resistance genes, including *Vibrio, Citrobacter, Aeromonas* and *Kluyvera*. Apart from *Vibrio*, these genera are not well-represented in the collection. Greater surveillance of these organisms could, as it has done for the other priority organisms, reveal an increasingly resistant trend and stimulate research, essential for the design of rational AMR control strategies.

Further to examining the count of resistance genes in discrete genomes, we have predicted how many classes of antimicrobials the genes within a genome confers resistance to. We find 35% of genomes (211,101/602,406) contain resistance to at least 3 classes of antimicrobials and have been defined here to be multi-class resistant (MCR). For a species to be described as MCR (red in Figure 3), at least half of the genomes from this species must be MCR (note this was only calculated for those species with at least 10 representatives). 37 species were classed as MCR. The WHO priority pathogens are well represented, though for *S. enterica* and *E. coli*, despite having some genomes conferring resistance to up to 12 and 14 different classes of antimicrobials respectively, the majority of samples are not MCR, though many may contain mutational resistance to antimicrobials such as fluoroquinolones. At the other end of the spectrum is *Enterobacter bugandensis*, where all 10 samples (from 3 different projects) contain genes conferring resistance to 8 classes of antimicrobials. *E. bugandensis* was only identified in 2016 and was associated with neonatal sepsis (31). The species *K. intermedia* and *V. cholerae*, in addition to possessing overall high numbers of AMR genes (Figure 3A), were also MCR. So too were the emerging opportunistic human pathogens *Raoultella planticola* (32) and *Corynebacterium striatum* (33) as well as the zoonotic pathogen *Histophilus somni* (34) and *M. tuberculosis*. However, the level of resistance in *M. tuberculosis* is likely to be underestimated as the main mechanism of resistance is through mutation (35) and so are not considered here.

**Figure 3.**
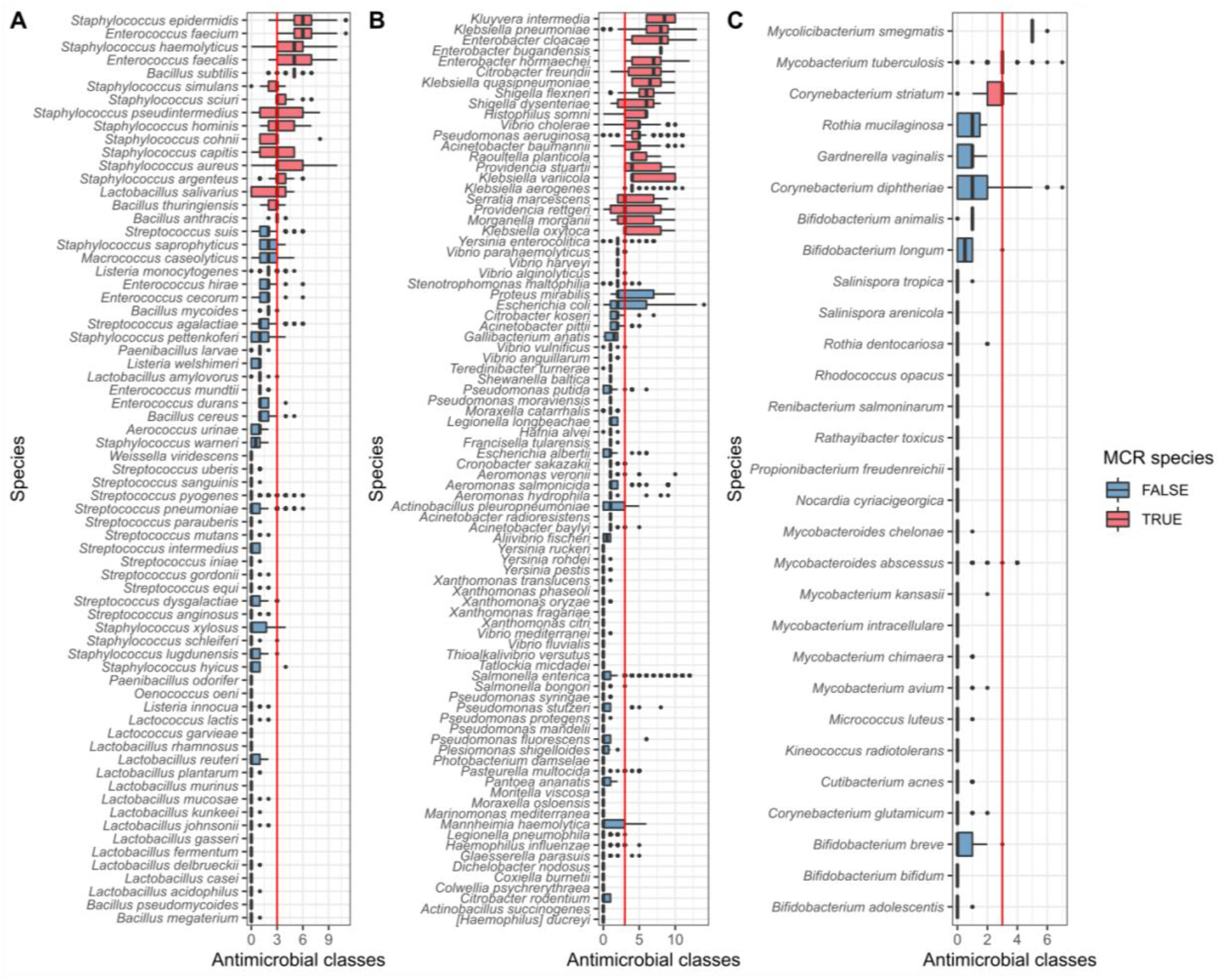
Predicted antimicrobial resistance profiles of species from A) Bacilli, B) Gammaproteobacteria and C) Actinobacteria, showing the number of predicted antimicrobial classes each isolate is resistant to, based on genetic profile. The red line indicates the threshold for MCR (predicted resistance to three classes of antimicrobials or more). Species are classed as MCR (red in figure) if at least 50% of the assemblies are MCR. Species included have at least 10 assemblies.

## DISCUSSION

Bacteria are a vast, diverse and ancient family of single-celled organisms that dominate this planet. In our efforts to understand and categorise this most abundant life form, hundreds upon thousands of bacterial sequences are submitted yearly into sequence archives such as the ENA. In the last two decades and with the advent of cheap high throughput short read sequencing the trend has moved away from the submission of finished or draft genome assemblies to one where simply the raw reads are submitted to public archives. These data usually require substantial preprocessing before they are analysis-ready. This takes significant time, expertise and computational power to do. By uniformly processing the data present in the ENA in November of 2018, we have collated a set of 661,405 standardised assemblies.

The additional standard characterisation and quality control we have performed enables the data to be easily subsetted for the purposes of identifying all the assemblies of a particular species or sequence type, or to those containing a specific antimicrobial resistance gene. Furthermore, this dataset can be interrogated for a specific gene or mutation through the use of the COBS search-index, for a specific genome by use of the provided minHash index and glean estimations of genetic distances of genomes of interest using the pp-sketch index. These facilities hint at the power of this unified resource, allowing phylogenetic relationships between genomes to be quickly elucidated, and hypotheses rapidly tested. This resource will empower more scientists to harness the multitude of data in the ENA both for surveillance and public health projects, as well as to address questions of basic science.

The count of 2,336 species in this snapshot is well below the number of bacterial species in the taxonomic databases such as NCBI taxonomy (>20,000 species, https://www.ncbi.nlm.nih.gov/Taxonomy/Browser/www.tax.cgi?id=2) and GTB (>30,000 species, https://gtdb.ecogenomic.org/). Some of the sequence diversity within the snapshot may have been missed due to limitations of the Kraken database used for taxonomic assignment and abundance estimation, a research project in its own right. For a small proportion of the assemblies (6.1%), a major species could not be assigned with high confidence, despite being shown by CheckM to contain little or no contamination, indicating that there was not a good match for it in the database (see Methods, Supplementary Figure 1D). The inclusion of genomes originating from metagenomic sequences from different sources (*e.g*. gut, skin, soil, ocean) would likely improve the overall species diversity but the methods of assembly and analysis are very different to those used here.

Many of the sequenced genomes could be defined as MCR based on the carriage of AMR genes. While we observe many occurrences of antimicrobial resistance mechanisms in the 661K assemblies, both in the organisms which are already know to be problematic (species outlined on the WHO priority pathogens list) and in newly emerged threats (such as *E. bugandensis, C. striatum* and *R. planticola*), it is difficult to estimate how well these reflect the true prevalence of resistance in a given species. This is due to many projects implementing pre-selection steps with only the antimicrobial resistant strains being then sequenced (36–38). This intrinsically biases the archive, preventing prevalence estimations. It also limits the power to track the origins of accessory genes and consequently the species interactions that can be inferred from this. Ideally, strategies to sequence a wider variety of species, including susceptible isolates, from diverse environments and global locations must be implemented before the dynamics of gene flow can be accurately studied.

The uniform resource of 661K bacterial assemblies that we present here removes several technical barriers to harnessing the wealth of public data stored in the ENA, enabling a broader community to access and leverage this data for their research. We envisage this to be a valuable resource which can provide the substrate for a wide range of future studies. Nevertheless, it is intrinsically limited through the nature of our scientific practice, by the diversity of sequences it holds. Rather, the current composition highlights the influences of the past quarter of century of funding and scientific focus. The enormous contribution of just a few projects shows that even the drive and focus of individual groups has influenced our view of recent bacterial diversity. Sampling and sequencing strategies must change if we want to reveal the bacterial tree of life.

## METHODS

### Download of reads, assembly and characterisation of genomes

The bacterial WGS datasets in the ENA as of the 26-11-18 were downloaded and assembled as a part of an assembly pipeline (https://github.com/iqbal-lab-org/assemble-all-ena) (39, 40). Only paired-end reads were included and those where the instrument platform was ‘PACBIO_SMRT’ or ‘OXFORD_NANOPORE’ were excluded. In addition, those with a library source of ‘METAGENOMIC’ and ‘TRANSCRIPTOMIC’ were also ignored. Available metadata and appropriate reads were downloaded and if multiple read sets were available they were appended together. Reads were assembled using Shovill v1.0.4 (T. Seeman, https://github.com/tseemann/shovill) with default options. Shovill uses SPAdes (v3.12.0) (11) for assembly, and includes some additional pre- and post-processing steps that utilise Lighter (41), FLASH (42), Trimmomatic (43), SAMtools (44), BWA-MEM (45, 46), seqtk (https://github.com/lh3/seqtk), Pilon (47) and samclip (https://github.com/tseemann/samclip), to speed up the assembly and to correct minor assembly errors. 664,877 assemblies were produced by this pipeline.

Separate from the assembly pipeline, Kraken v2.0.8-beta (9) was run on the read fastq files using the Kraken2-microbial database (2018, 30GB) and the resulting taxonomy labels assigned by Kraken were analysed by Bracken v2.5 (10) to estimate the species abundance within each set of reads. From the assemblies, contigs of less than 200 bp were removed using the script available at https://github.com/sanger-pathogens/Fastaq and contigs of *k*-mer depth less than 10 were noted, but not removed. Quast version 5.0.2 (12) was used to summarise assembly statistics and CheckM v1.1.2 (13) using the “--reduced_tree” flag was used for estimations of completeness and contamination of an assembly. Assemblies with a genome length of less than 100 Kb or longer than 15 Mb were removed (3,472 assemblies), leaving 661,405 assemblies. A minHash sketch of each assembly (“-n 5000”) was produced using sourmash v3.5.0 (14). A searchable k-mer database of the 661K assemblies was constructed by COBS (checkout 7c030bb) using “compact-construct” with default options (8). Core and accessory distances were calculated between the assemblies using poppunk_sketch v1.5.1 with default options except “--k-step 3” (15). MLST was determined where possible using mlst v2.19.0 (Seeman, T. mlst, https://github.com/tseemann/mlst), *E. coli* phylotype determined using clermonTyping version 1.4.1 (15) and *Salmonella* were serotyped using SeqSero2_package.py v1.1.1 (16). Plasmid replicons were detected using Abricate v1.0.1 (Seeman, T. abricate, https://github.com/tseemann/abricate) with the plasmidfinder 2020-May-7 database (17) and AMR, heavy metal and virulence genes were detected using AMRFinderPlus v3.6.15 (18), with standard thresholds of minimum identity (curated cut-off if it exists and 0.9 otherwise) and default coverage of 0.5. All figures were generated in R using ggplot2 (19) and where required were edited manually using Inkscape 2 v0.92.

### Taxid lineage, species comparison and adjustment species abundance

The taxid lineage of the major bracken species was acquired by NCBITaxa (20). Where the major species from the Bracken analysis belonged to either of the *M. tuberculosis* complex or *B. cereus* s.l. complex or was identified as a *Shigella* sp. or an *E. coli*, the remainder of the read assignments were examined to see if they belonged to other members of that complex. If they were members, their assigned percentage was added to that of the major species.

### High quality assemblies

Filtering was applied using the reports generated by Quast and CheckM analysis for each genome. The high quality assemblies met the requirements of: less than 2,000 contigs, a genome length that is within the acceptable range for that species (50%-150% of the expected length) (ftp://ftp.ncbi.nlm.nih.gov/genomes/ASSEMBLY_REPORTS/species_genome_size.txt.gz, 27^th^ August, 2020), or is unknown, a N50 of greater than 5,000, a completeness score of at least 90 and a contamination score of less than or equal to 5. In total, 639,981 assemblies met these requirements.

### Multi-class resistance

Multi-class resistance (MCR) was defined as containing genes conferring resistance to at least 3 classes of antimicrobial (antimicrobial classes were extracted from the AMRFinderPlus output). Only species with at least 10 samples were included and a species was classed as MCR if at least 50% of individual assemblies were MCR.

## DATA AND CODE AVAILABILITY

The 661,405 assemblies as well as the COBS, minHash and pp_sketch indices are available: ftp.ebi.ac.uk/pub/databases/ENA2018-bacteria-661k.

The pipeline used for download and assembly of reads from the ENA https://github.com/iqbal-lab-org/assemble-all-ena.

Additional metadata and characterisation files deposited in figshare (https://dx.doi.org/10.6084/m9.figshare.14061752):

- Full metadata downloaded from the ENA for each assembly in json form (Json1_ENA_metadata)
- Full QC and general characterisation including AMR gene and plasmid replicon detection, for each assembly in json form (Json2_QC_characterisation_amr_plasmid)
- Kraken/Bracken output including the top 50 species for each assembly (File1_full_krakenbracken)
- The taxid lineage of the major species determined using NCBITaxa (File2_taxid_lineage_661K)
- Summarised metadata from the ENA for each assembly (File3_metadata_661K)
- Summarised QC and general characterisation for each assembly (File4_QC_characterisation_661K)
- Summarised AMR genes, MCR status, plus genes, plasmid replicons for each assembly (File5_AMR_plasmids_661K)
- Presence/absence matrix of AMR genes in each assembly (File6_AMR_presenceabsence_661K)
- Class of each AMR gene extracted by AMRFinder (File7_gene_class_AMRFinder)
- Presence/absence matrix of plasmid replicons in each assembly ( File8_plasmidreplicons_presenceabsence_661K)

R notebooks used for analysis and figure generation have been deposited in figshare ():

- Code used to generate figures in the QC and filtering section (Rnotebook1_QC_filtering_section)
- Code used to generate figures in the Species breakdown section (Rnotebook2_species_breakdown_section)
- Code used to generate figures in the AMR section (Rnotebook3_AMR_section_figures)

## Author contributions

G.A.B., Z.I. and N.R.T. conceptualised the project. M.H. wrote the assembly pipeline which was run by G.A.B., M.H. and K.M.M. Species identification was performed by G.A.B. and B.T.F.A. G.A.B performed QC and characterisation of assemblies and with the help of G.H., analysed and visualised the results. The minHash and pp-sketch indexes were constructed by G.A.B. and the COBS index was constructed by L.L. and G.A.B. The manuscript was written by G.A.B, Z.I. and N.R.T. All authors read and approved the final manuscript.

## Funding

This work was supported by Wellcome (206194). G.A.B. was funded by an EMBL-EBI and Sanger ESPOD fellowship. M.H. was funded by a Wellcome Trust/Newton Fund-MRC Collaborative Award (200205) and an award from the Bill & Melinda Gates Foundation Trust (OPP1133541).

## Conflicts of interest

The authors declare no conflicts of interest.

## Acknowledgements

We thank Alexandre Almeida, Kate Mellor, Alyce Taylor-Brown and all other members of the Iqbal and Thomson research teams for their useful discussions and suggestions. We would also like to thank John Lees for his helpful guidance and support when creating the pp-sketch index of the 661K assemblies.

## Notes

### Competing Interest Statement

The authors have declared no competing interest.

