## Supplementary material for "Exploring bacterial diversity via a curated and searchable snapshot of archived DNA sequences"

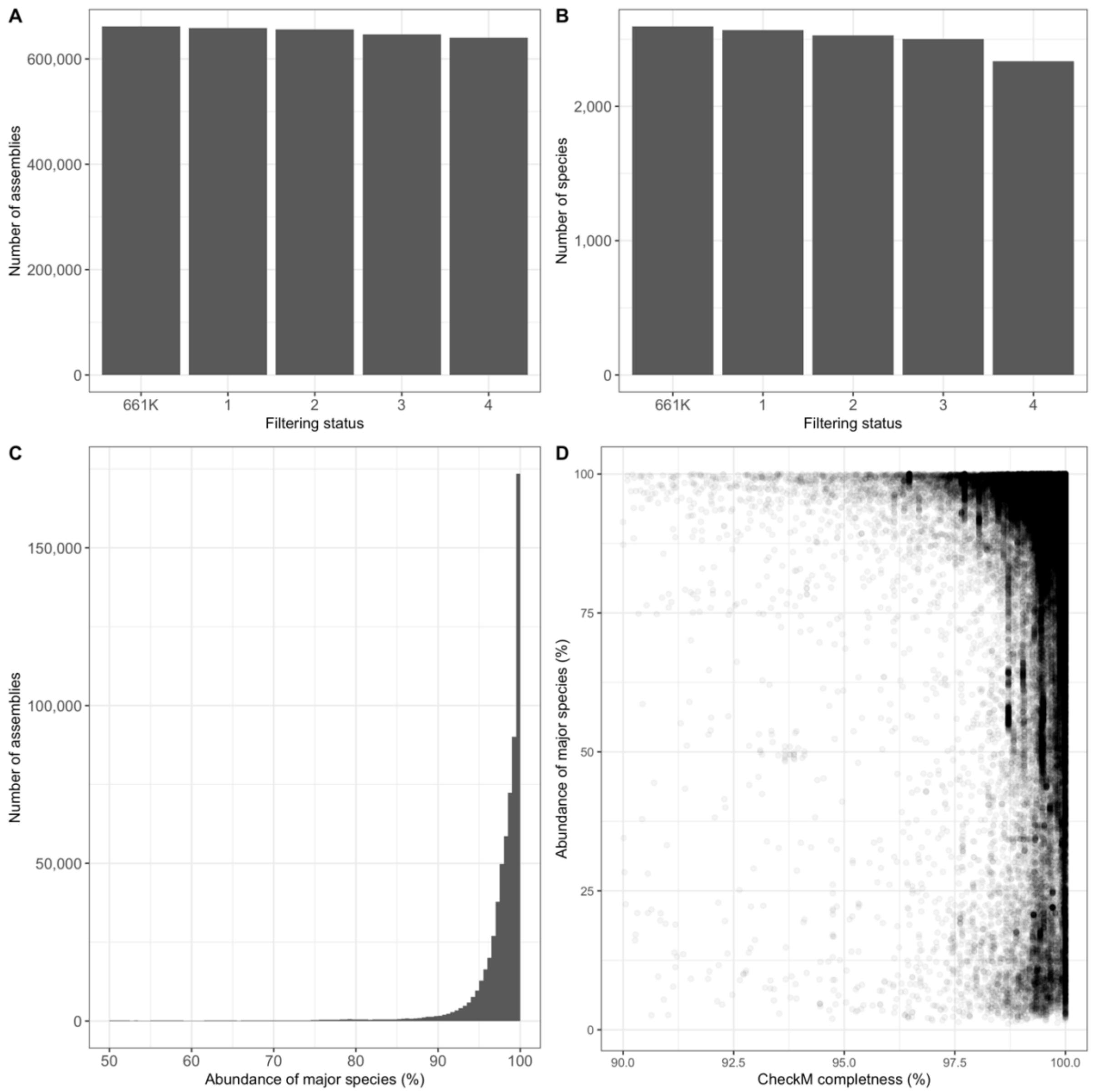


**Supplementary Figure 1.** QC of the 661,405 assemblies. Number of A) Assemblies and B) species remaining following each stage of filtering. Status 1; removal of genomes with > 2000 contigs, status 2; removal of genomes with an N50 < 5000, status 3; removal of genomes with length outside the range expected for that species (note: if expected range is not known, the assemblies are kept), status 4; assemblies with a completeness score =>90% and with a contamination score =< 5%. 639,981 assemblies passed the four levels of filtering and are the high-quality genomes. C) Distribution of high-quality assemblies (filtering status 4) with > 50% abundance of major species. 9,595 assemblies were below this threshold. D) Within-sample abundance of major species vs completeness of the high-quality assemblies. For C) and D) the abundance of major species is the adjusted abundance values (see methods).

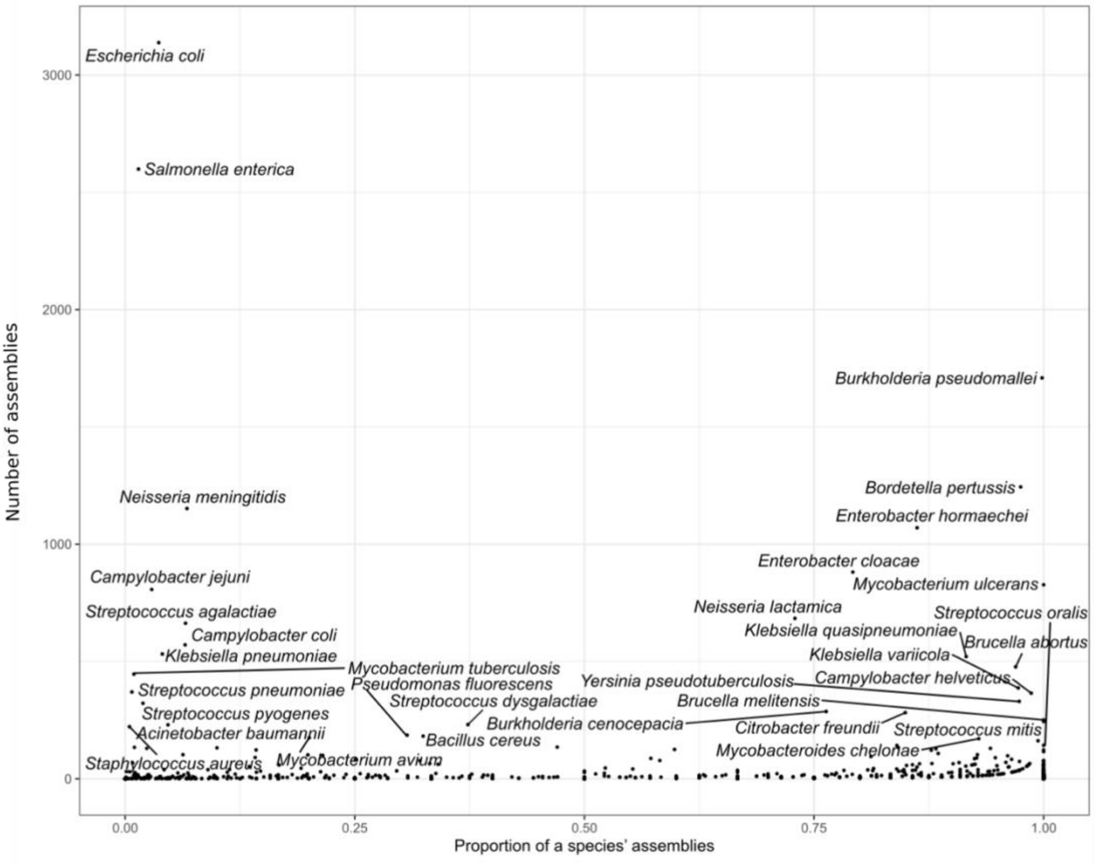


**Supplementary Figure 2.** High quality assemblies which have a major species abundance of less than 90%.

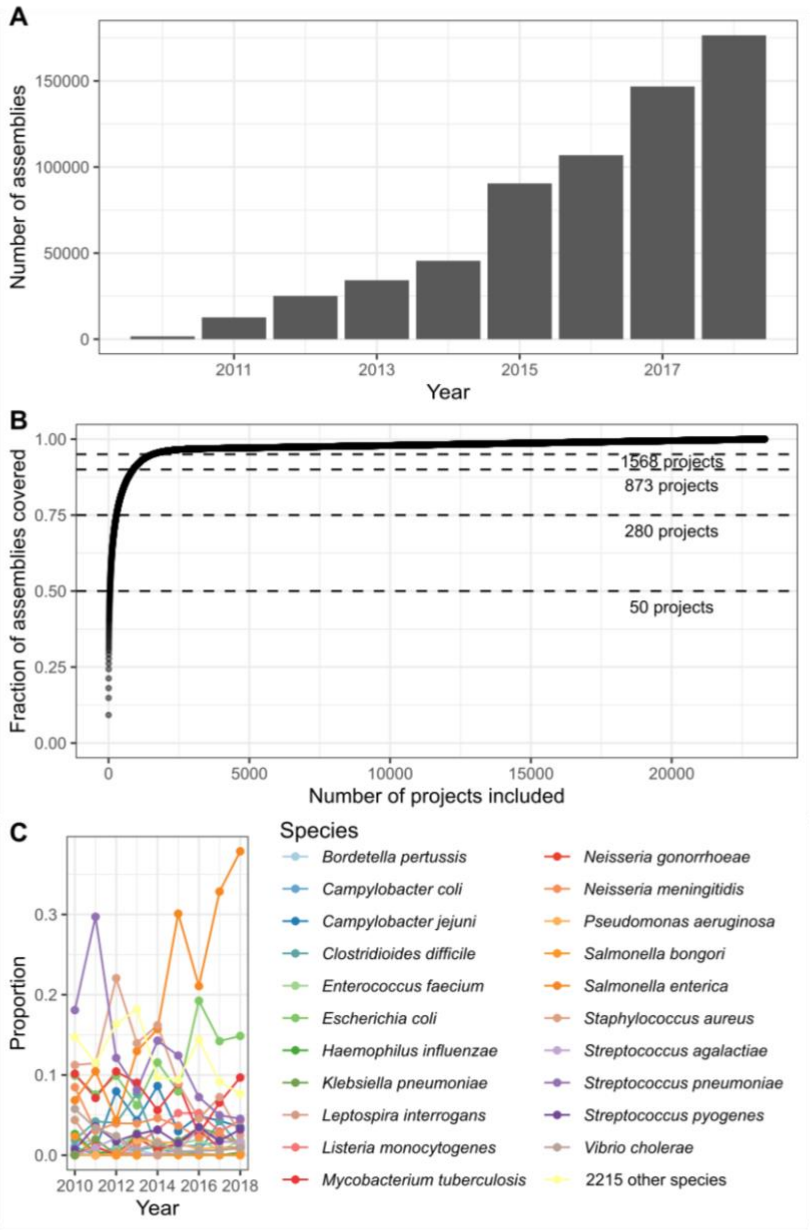


**Supplementary Figure 3.** Composition of the 639,981 high quality assemblies. A) Breakdown of assemblies by year first public in the ENA. B) Fraction of assemblies covered by accumulating projects. C) Tracking proportions of the top 10 bacterial species for a year.

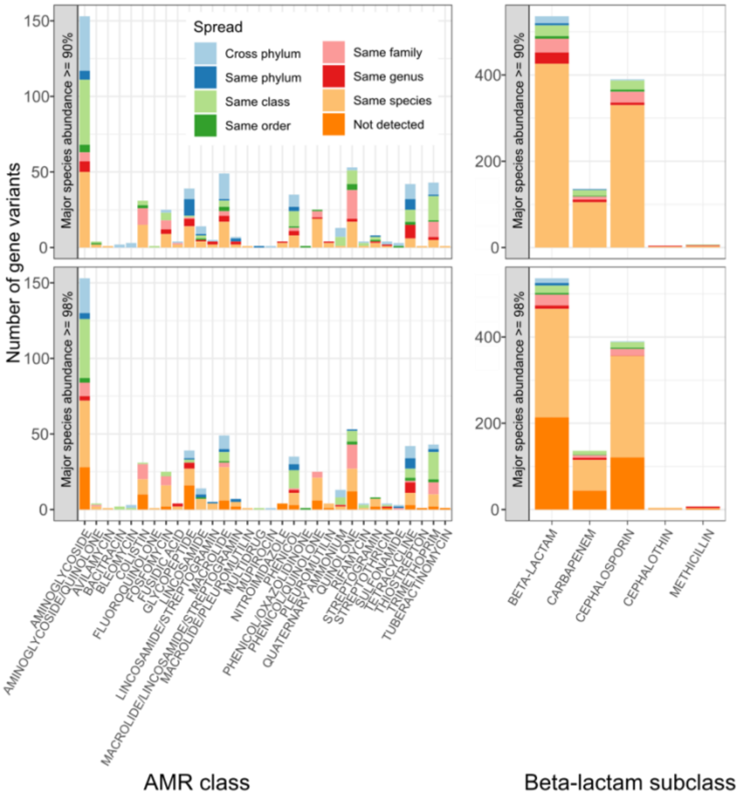


**Supplementary Figure 4.** Distribution of antimicrobial resistance gene alleles by antibiotic class. Gene variants are coloured by their level of spread, from being detected in genomes from different phyla to only found in a single species. The top graph includes genomes with the major species being >= 90% abundance and the lower graph is when this threshold was increased to >= 98% abundance.
